# Branching and oscillations in the epigenetic landscape of cell-fate determination

**DOI:** 10.1101/007831

**Authors:** Jomar Fajardo Rabajante, Ariel Lagdameo Babierra

**Affiliations:** Institute of Mathematical Sciences and Physics, University of the Philippines Los Baños, College, Laguna 4031 Philippines

**Keywords:** gene regulatory network, stem cells, pluripotency, synthetic biology, multistability, attractor

## Abstract

The well-known Waddington’s epigenetic landscape of cell-fate determination is not static but varies because of the dynamic gene regulation during development. However, existing mathematical models with few state variables and fixed parameters are inadequate in characterizing the temporal transformation of the landscape. Here we simulate a decision-switch model of gene regulation with more than two state variables and with time-varying repression among regulatory factors. We are able to demonstrate multi-lineage differentiation at different timescales that portrays the branching canals in Waddington’s illustration. We also present a repressilator-type system that activates suppressed genes via sustained oscillations in a flattened landscape, hence providing an alternative scheme for cellular reprogramming. The time-dependent parameters governed by gradient-based dynamics regulate cell differentiation, dedifferentiation and transdifferentiation. Our prediction integrates the theories of branching and structural oscillations in cell-fate determination, which reveals key temporal patterns of cell differentiation and associated diseases, such as cancer.

## Main text

Waddington’s epigenetic landscape illustrates the canalization in the cell differentiation and fate determination process [1–4]. The topography of Waddington’s illustration represents the developmental pathways of tissues formed from totipotent and pluripotent cells to terminally-differentiated specialized cells (Fig. 1a). Various theoretical studies have quantified Waddington’s epigenetic landscape and are able to predict bistability in gene regulatory networks (GRNs) [5–9]. However, many of the mathematical models only consider at most two regulatory factors, and focus on static epigenetic landscape represented by fixed parameter values. In reality, the topography of Waddington’s illustration is dynamic and involves many regulatory factors (high-dimensional [10,11]). The parameters that represent gene regulation are indeed changing during the development of an organism [10,12–17]. Mathematical models with two regulatory factors and fixed parameter values only describe a particular temporal scenario in cell differentiation.

**Figure 1.**
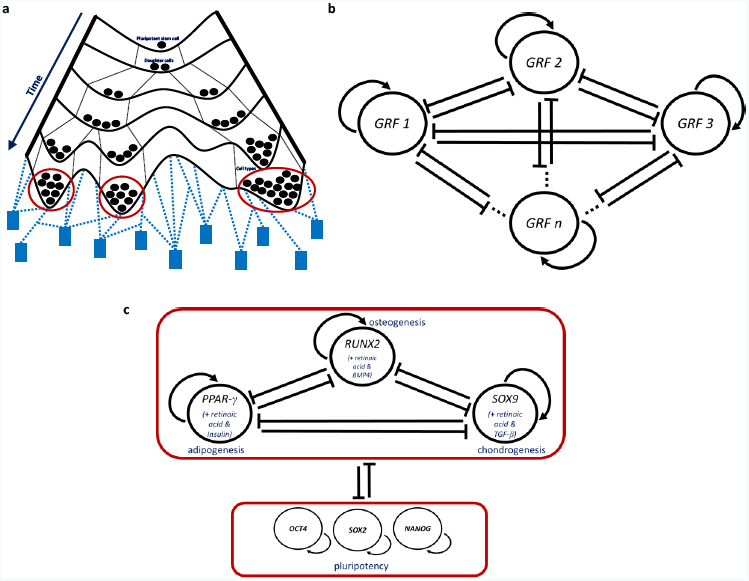
A sketch of the epigenetic landscape, and a gene regulatory network (GRN) for cell-fate determination. (**a**) Epigenetic landscape adapted from Waddington’s illustration [1]. The branching canals depict the various cell lineages towards different fates (cell types/phenotypes). The cell fates are illustrated as valleys and traditionally represented as mathematical attractors (see Supplementary Figs. 1 and 2). The differentiating cells, illustrated as balls, trail a chosen canal towards a specific valley. The canal is chosen based on the landscape’s potential (similar to gravitational potential) such that the steeper pathway and deeper valley are preferred [8]. The canals are separated by ridges that restrain cells to switch lineages. The blue pegs (GRFs) and strings attached to the pegs (GRF-GRF interaction) alter the height of ridges and depth of valleys. The height of ridges and depth of valleys vary through time and affect the route of the differentiating cells. The transformation in the phase portrait of the dynamical system depicts the changes in the topography of the epigenetic landscape (Supplementary Fig. 2). (**b**) A minimal GRN that characterizes decision switches in cell-fate determination. We suppose that a node represents a GRF involved in expressing a specific cell type, such as master switch genes, transcription factors, or coarsegrained modules of a larger GRN that can be simplified as one node. Each node has auto-activation as represented by the arrows; while the interaction links among GRFs is of repressive behavior (represented by the bars). The repressive behavior of GRFs denotes that a mature cell only expresses one phenotype and hinders the expression of multiple phenotypes. Strength of repression is not necessarily reciprocal and a one-way repression is possible. The auto-activation and repression can be direct or indirect. Examples of GRNs of this type are discussed in various literatures [10,37,43,75]. (**c**) A course-grained mesenchymal stromal transcription network. The upper module consists of lineage-specifying master genes, and the lower module consists of pluripotency factors that maintain the identity of stem cells. Activation of the upper module and inactivation of the lower module drive lineage commitment towards adipogenesis, osteogenesis or chondrogenesis.

The mechanisms that regulate gene expression, such as kinetics of gene regulatory factors (GRFs) and the structure of GRF-GRF interaction, influence the outcome of cell-fate determination [7,10,12,15–18]. Waddington observed that changes in these mechanisms could alter the epigenetic landscape leading to cell-lineage switching [1]. The changes in the GRF-GRF interaction do not necessarily entail mutations but can be due to normal processes. In mathematical point-of-view, the variations in gene regulation can be represented by modifications in the parameter values of the quantitative models. Bifurcation analyses of existing models have been done [5,7,12,16,17], but most of them do not provide elaborate illustrations of cells trailing the dynamic pathways governed by more than two GRFs. Here we present numerical illustrations of cells trailing different epigenetic routes such that the pathways transform due to changes in the strength of repressive interaction among multiple GRFs (see Fig. 1b and Box 1 for the mathematical model). The GRFs in the model (Fig. 1b) have mutual repression because a mature cell expresses only one phenotype and constrains the expression of the other phenotypes.

An example of GRN with mutual repression is the coarse-grained mesenchymal transcription network shown in Figure 1c. The upper module consists of lineage-specifying master genes, namely, PPAR-γ, RUNX2 and SOX9 [18]. Auto-activation is a common property of master genes [10]. In mouse embryonic stem cells, the up-regulation of PPAR-γ by retinoic acid and insulin, while inhibiting the growth of RUNX2 and SOX9, steers adipogenesis (fat formation). The up-regulation of RUNX2 by retinoic acid and BMP4 represses PPAR-γ and SOX9 and drives osteogenesis (bone formation). On the other hand, retinoic acid and TGF-β up-regulate SOX9 leading to chondrogenesis (cartilage formation) and to the inhibition of PPAR-γ and RUNX2 [18]. The activation of the upper module (Fig. 1c) represents cell differentiation towards specific lineages and is hindered by the module of pluripotency factors (lower module in Fig. 1c).

A desired cell fate can be a cell type/phenotype that is essential for proper normal development, or desired cell type during cellular engineering. Our main assumption is that gradient-based optimization governs the transformations of the pathways leading to the desired cell fate, following the theory that differentiating cells choose the steeper pathways (canals) in the epigenetic landscape. This assumption assures that the cells trail the nearby steepest pathways shaped by the time-varying antagonistic interaction among the GRFs (see Methods). The gradient-based method can be considered as a cell-fate induction strategy such that the cells move from a pluripotent state, which has higher network entropy, towards differentiated state with lesser network entropy [19]. This dynamic GRF-GRF interaction demonstrates the cascade of branching canals in Waddington’s illustration. It also describes cell plasticity by allowing cell-lineage switching by transdifferentiation and dedifferentiation.

#### Box 1: *Mathematical model*

We consider a minimal gene regulatory network (GRN) of the form shown in Figure 1b, where the number of nodes (*n*) is arbitrary. This GRN is a representation of the network that characterizes decision switches in cell-fate determination [10,37,43,50,75]. For simplicity, we draw the network in such a way that each node represents a gene regulatory factor (GRF) involved in expressing a specific cell type/phenotype [10,13]. A node in this GRN represents either a specific GRF, or a coarse-grained subnetwork of multiple factors that can be treated as single GRF. An example of a GRN in the form shown in Figure 1b is the coarse-grained mesenchymal transcription network with PPAR-γ, RUNX2 and SOX9 as nodes (Fig. 1c) [18].

One of the simple models that describe the qualitative dynamics of the GRN with multiple GRFs is the following system of differential equations:

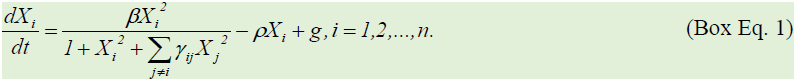

The state variable *X*_*i*_ represents the strength/concentration of the GRF involved in expressing the gene towards the *i*-th cell type. The parameters *ß>0, p>0, g*≥*0* and *γ*_*ij*_≥*0* are the efficiency of GRF in expressing the corresponding gene, degradation rate, basal constitutive growth rate, and time-varying interaction coefficient associated with the inhibition of *X*_*i*_ by *X*_*j*_, respectively. The kinetics of GRF auto-activation is a sigmoidal increasing function that is negatively influenced by the strength of the other GRFs. This model is originally proposed by Cinquin and Demongeot [10,13] but their model has symmetric parameters unlike Box Eq. 1 which admits asymmetric repression coefficient *γ*_*ij*_.

We assume that the goal of gene regulation is to maximize the strength of GRFs so that the outcome is moving towards the steepest canal and possibly towards the deepest valley in the epigenetic landscape, which is expected in cell-fate determination. Here we apply a gradient-based optimization method to the time-varying repression strength *γ*_*ij*_ to drive cell-fate induction in the direction of the nearby steep canal (see Methods). This method is governed by a timescale factor that declines through time [5,12].

The equilibrium points and sustained oscillations of this model lie on the hyperspace 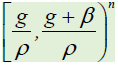. The model has at most *3*^*n*^ equilibrium points. See [35] and Methods for additional information about the mathematical model.

*Basic model of gene expression.* In order to understand the dynamics of gene regulation with multiple GRFs, it is helpful to recall the basic models of gene expression involving a single GRF. One of the basic models can be written as

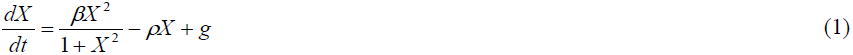

where *X* represents the strength/concentration of the GRF, *β>0* is the gene expression efficiency, *ρ>0* is the degradation rate, and *g≥0* is the basal constitutive growth rate of the GRF. The sigmoid growth curve and the decay line (Fig. 2) are respectively defined by

**Figure 2.**
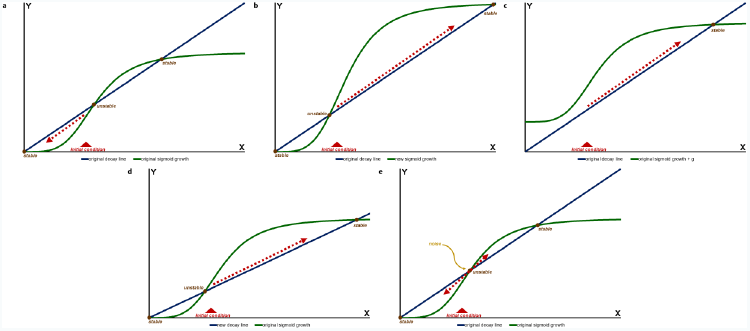
Graphical analysis of the basic model of gene expression. The intersections of the sigmoid growth curve and the decay line are equilibrium points. Suppose we have a fixed initial condition. (**a**) The initial condition prompts convergence to low-valued stable equilibrium point (down-regulation of gene expression). (**b**) Increasing the value of the gene expression efficiency *β* changes the geometrical shape of the sigmoid growth curve, which now results in the convergence to high-valued stable equilibrium point (up-regulation of gene expression). (**c**) Adding external stimulation that augments the effect of the basal constitutive growth rate *g* also changes the geometrical shape of the sigmoid growth curve, which results in the convergence to high-valued stable equilibrium point (up-regulation of gene expression). Notice that it is possible to eliminate the existence of low-valued stable equilibrium point such that only the high-valued equilibrium remains. (**d**) Decreasing the slope of the decay line can also lead to high-valued stable equilibrium point (up-regulation of gene expression). (**e**) Without parameter modification, stochastic fluctuations regulate activation and inactivation of genes. Stochastic noise can drive cell-lineage switching.

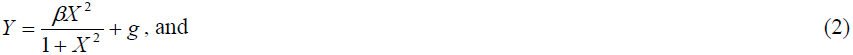

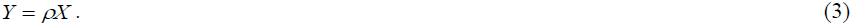

Investigating the effect of dynamic model parameters provides insight on the up-regulation and down-regulation of a gene. For example, it is possible to reverse the fate of an initial condition that leads to the down-regulation of a gene (convergence to low-valued stable equilibrium point; see Fig. 2a) by increasing the value of gene expression efficiency *β* (Fig. 2b) or by adding external stimulation that augments the effect of the basal constitutive growth rate *g* (Fig. 2c). Enhancement of the maximal growth rate *β+g,* such as by introducing external stimulus, is a direct technique in activating genes [20–24]. Another technique to drive gene activation is the reduction in the degradation rate *ρ* (Fig. 2d) [25,26]. Extrinsic and intrinsic stochastic noise also plays significant role in the up-regulation and down-regulation of gene expression (e.g., Fig. 2e), as discoursed in various theoretical and experimental studies [18,27–33].

Nevertheless, this basic model of gene expression is often not enough to explain the complexity of cell-fate determination. To advance the theory of gene regulation, it is necessary to include many GRFs because there are key dynamics that are not conceivable in a model with only one variable. The canalization and oscillations in Waddington’s epigenetic landscape are observable in an ordinary differential equation model with at least two or with at least three state variables, respectively. The oscillation-induced transdifferentiation and dedifferentiation discussed in this study are only possible with GRNs involving three or more GRFs.

## Results

We observe two significant dynamics in our simulations. The first one is multilineage differentiation via sequentially branching developmental pathways. This sequential branching portrays the canalization in Waddington’s landscape at different timescales. The pathways trailed by the differentiating cells depend not only on the structure of the GRN and parameter values but also on the initial condition (see Supplementary Fig. 1). The second one is flattening of the epigenetic landscape which eliminates the deep valleys, resulting in sustained oscillations. The GRN that generates the oscillations (oscillator) can boost the strength of a suppressed GRF in partially or terminally-differentiated cells. This oscillatory behavior is structural in nature (i.e., based on the structure of the GRN) and not due to noise nor delay. Therefore, we are able to integrate into a single theory the theories of branching and structural oscillations in cell-fate determination.

Mathematically, the sequential branching in the epigenetic landscape towards different cell types represents convergence to one of the equilibrium points (Supplementary Fig. 1). The variations in the interaction coefficients drive changes in the topography of the landscape (Supplementary Fig. 2) and eventually stabilize at different timescales, hence sequential branching become possible (refer to the timescale factor in Methods). The pathway bifurcates from one branch (a primed state) to multiple branches (lineages), which may further have sub-branches that culminate in branch endpoints (cell-fates); see Figures 3a, 3c-3e and Supplementary Figure 21. The final structure of the GRN, which is determined by the parameters stabilized at different timescales, dictates the number and location of the branch endpoints (Figs. 3a-3e; see Supplementary Fig. 21 for a system with 10 GRFs). Generally, in order for sequential branching to arise, the initial structure of the GRN and parameter values should allow bistability or multistability which is a property of multipotent and pluripotent cells (Supplementary Fig. 1). The network entropy and the quasi-potential of the landscape are lower in populations of differentiated cells compared to the undifferentiated state as expected (e.g., Fig. 3f and see the quasi-potential axis in Figs. 3a and 3b; refer to Methods for the description of network entropy and quasi-potential). In addition, the pathways with two endpoints that are distant from each other are usually more robust against stochastic noise compared to the pathways with many endpoints (Supplementary Figs. 3, 6, 8 and 10). This implies that cells trailing the pathways with more endpoints can be candidates for stochastic direct transdifferentiation.

**Figure 3.**
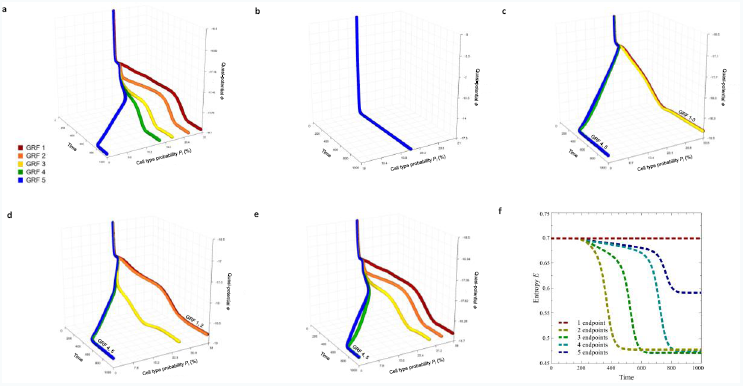
Illustrations of cells trailing the branching pathways in the epigenetic landscape. (only the deterministic path is shown; see Supplementary Figs. 3, 5, 6, 8 and 10 for stochastic simulations and for the initial condition and parameter values used). Note that the branch endpoints are the coordinates of an equilibrium point. The different numbers of endpoints are due to the different timescale factors used. (**a**) Five different endpoints. (**b**) No branching and only one endpoint (graphs for GRF 1-5 are superimposed to each other). The state with equal cell type probabilities represent multipotent or pluripotent cells (undifferentiated state). (**c-e**) Two, three and four endpoints, respectively. (**f**) Time series of the network entropy levels for the system in Figures 3a to 3e. The network entropy decreases as cells differentiate.

There are cases where a stable equilibrium point vanishes and a stable limit cycle emerges, especially when the repressive reciprocal interaction among GRFs is asymmetric (Fig. 4a). This attracting limit cycle is generated by an oscillator that perpetually up-regulates suppressed GRFs and down-regulates dominant GRFs, resulting in fluctuating kinetics (Fig. 4b). One example of an oscillator is a repressilator-type network (Fig. 4c), which is similar but not exactly identical to the repressilator proposed by Elowitz and Leibler [34]. In this repressilator-type network, the strength of repression in one loop is stronger than the strength of repression in the reverse loop (asymmetric interaction). The sustained oscillations generated by this repressilator-type network can arise in a GRN with three or more nodes (odd or even number of GRFs; e.g., Figs. 5a and 5b). The dynamics of this oscillator can be illustrated by an epigenetic landscape with flattened topography, that is, there are no deep valleys in the route of the differentiating cells, and the cells are continually sliding in zigzag canals without endpoint.

**Figure 4.**
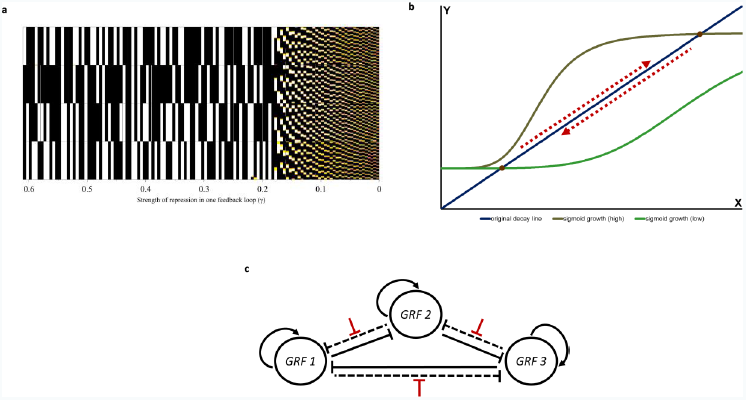
Oscillating pathways in the epigenetic landscape generated by repressilator-type networks. (**a**) Hopf bifurcation colormap diagram showing the birth of limit cycle. Suppose *n=3*, *a*_*12*_*=a*_*23*_*=a*_*31*_*=3*, *a*_*13*_*=a*_*21*_*=a*_*32*_*=γ* and *g=0.05; ε*_*1*_*=ε*_*2*_*=ε*_*3*_*=0*. Decreasing the value of γ creates a GRN such that one repression loop is stronger than the reverse loop. Pure black and pure white scale bars characterize equilibria. Colored scale bars characterize oscillations. (**b**) With more than one GRF (specifically, *n≥3*), the sigmoid growth curve continuously changes due to the asymmetric (negative feedback) repression loop. This results in the disappearance of equilibrium points and in the appearance of sustained oscillations. The oscillating solution to the differential equation model hovers between the high-valued and low-valued intersections of the decay line and the changing sigmoid growth curve (here, the intersections do not necessarily denote equilibrium points). Note that the basic model of gene expression (*n=1*) cannot demonstrate this type of qualitative dynamics. (**c**) A repressilator-type network with a strong asymmetric (negative feedback) repression loop, *n=3* (this can be extended for any *n≥3*). The strengths of repression in one loop (solid long bars) are stronger than the reverse loop (long broken bars). The short red bars represent inhibition of repression. Note that in our model, only three GRFs are needed to form a negative feedback repression loop that generates sustained oscillations. This repressilator-type network has auto-activation unlike the repressilator by Elowitz and Leibler [34].

The oscillations drive the strengths of the GRFs to have alternating positive and negative rates, whereas the gradient-based optimization method forces the dynamics towards positive rates only. Thus, oscillatory behavior is not optimal in the sense of differentiation towards cell types located at deep valleys in the landscape. We then expect that gradient-based dynamics that are persistent for some period result in damped oscillations (Figs. 5c and 5d). There are cases where the damped oscillations illustrate multipotency (or pluripotency depending on the GRN) which is represented by the equal probabilities of differentiating towards all the considered cell types (Fig. 5c). In some cases, the interaction coefficients vary with different timescales, resulting in partial differentiation and sometimes in the reversal of the status of the initially dominant GRF (Fig. 5d). Note that if the dampening of the oscillations is fast, the initial oscillations can be unnoticeable yet can still activate suppressed genes (Supplementary Fig. 19).

## Discussion

Various studies have attempted to model the cell differentiation process, but there are still more to uncover in epigenetics. Further theoretical prediction and experimental validation are needed to fully explain cell-fate determination and reprogramming. Varying the efficiency of GRF in expressing a gene (*β*), the degradation rate (*ρ*), or the constitutive growth rate (*g*) is a straightforward technique in stimulating the activation or deactivation of a GRF and its corresponding gene [6,7,10,18,35]. However, regulating the repression strength of GRFs (*γ*_*ij*_) has not been explored, and we have shown numerical illustrations where variations in this GRF-GRF repression affect the qualitative behavior of the cell differentiation system (Figs. 3 and 5). We are able to replicate Waddington’s model using a single set of equations with many GRFs involved. A GRN with only two nodes generally cannot describe the sequential bifurcation of canals and the oscillations in cell-fate determination. The different timescales involved in gene interaction influence the outcome of cellular regulation [5,36–39].

**Figure 5.**
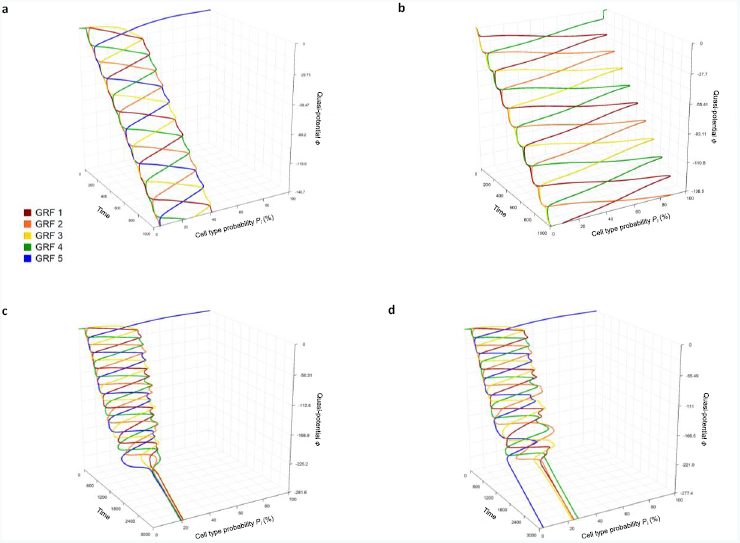
Examples of oscillating pathways. There are no deep valleys only continuous zigzag canals (see Supplementary Figs. 13-15 and 17 for the stochastic simulations and for the parameter values used). (**a**) *n=5*; *X*_*5*_ is initially silenced. (**b**) *n=4*; *X*_*4*_ is initially silenced. (**c**) Damped oscillations towards multipotency or pluripotency. The rates of decline of the timescale factors are all equal to *0.001*. (**d**) Damped oscillations resulting in partial differentiation and reversal of dominant GRF. The rates of decline of the timescale factors are not all equal. The initial dominant regulatory factor is GRF 5 but eventually becomes inferior as oscillations dampen.

One of the aims of this study is to spur more discussions on non-equilibrium dynamics and oscillations arising from asymmetric systems with multiple GRFs, which can broaden our understanding about the mechanisms of gene regulation. Reversal of the route from differentiated state to pluripotency is previously thought to be impossible but now many dedifferentiation techniques have been proposed and tested by experiments [9,40,41]. We propose another alternative technique for cellular reprogramming, which is by rewiring the GRN to have a repressilator-type network, possibly with the aid of external input and stochastic noise [18,42]. Exogenous inhibitors can be introduced to weaken the repression in one loop (Fig. 4c). Our numerical predictions can help design cellular engineering strategies for generating induced multipotent stem cells (or pluripotent stem cells depending on the GRN [18,43,44]) using the oscillations that can activate silenced genes. Several studies have discussed that oscillating GRF expression is indeed an attribute of progenitor cells [45–47], thus supporting our claim. However, note that in reprogramming back to pluripotency, we also need to assure activation of defined factors in the core pluripotency circuitry, such as *Oct4*, *Sox2* and *Nanog* [18,40].

The oscillator motifs (e.g., repressilators) which are part of a larger GRN contribute to the fluctuations observed in gene regulation dynamics. In fact, there are many types of oscillators [48–50]. The oscillations generated by various oscillator motifs when combined may produce fluctuations similar to stochastic noise. However, note that the combined large and small oscillations are sometimes not entirely random noise, especially when the detected fluctuations are part of the GRN structure and not coming from stochastic sources [29,51]. There are theoretical perspectives that interpret cell-fate determination as a chaos-driven process [52]. Another perspective is that the oscillator motif of a larger GRN can be used for artificial transdifferentiation by generating oscillations that can prompt cell-lineage switching, similar to what stochastic fluctuations can do [29,53]. In fact, direct transdifferentiation between related cell types branching from one lineage can be more straightforward compared to dedifferentiation to pluripotency [54].

From our simulations, we formulate some conjectures. (**i**) The dedifferentiation caused by abnormal oscillators (e.g., aberrant repressilator-type network) play a role in the existence of cancer stem cells and mutator phenotype [30,55–58]. Abnormal changes in the structure of GRF-GRF interaction, such as abnormal timescale factor and abnormal weakening of repression links, can lead to disease. Indeed, partially reprogrammed cells and excessive plasticity can cause cancer [30,54,59,60]. Moreover, it is also possible that these oscillators play a part in epigenomic reprogramming and influence transgenerational epigenetic inheritance [61]. Abnormalities in the GRF-GRF interaction could be passed-on to offspring. (**ii**) We can reprogram cells back to pluripotency by regulating the wiring of the GRN. This implies that there are no unique reprogramming factors, and we can reprogram cells using any regulatory factor as long as it can lower the “gravity” of the epigenetic landscape [54]. In this paper, we have shown theoretical predictions of novel developments in the theory of gene regulatory networks. One experimental approach to demonstrate our numerical predictions is by employing synthetic biology techniques [33,34].

In reality, the temporal transformation in the epigenetic landscape is due to multiple intrinsic and extrinsic factors. Here we only consider changes in the GRF-GRF interaction coefficient *γ*_*ij*_ but we should not disregard that gene regulation consists of the interplay among many factors and processes. For example, GRF-GRF interaction can be regulated not only through *γ*_*ij*_ but also through the modifications in the maximal growth rate *(β+g)* or through the degradation rate *(ρ)* [6,7,10,18,35]. Increasing the maximal growth rate or decreasing the degradation rate of a certain GRF enhances the steady state strength of the GRF, which in turn intensifies the repression of the other GRFs. Furthermore, the dynamics observed from empirical data combine the various effect of many parameters. For example, a decline in the strength of a GRF suggests various possible reasons, such as due to an increased degradation rate or due to an increased repression by an antagonist GRF. Hence, we need to interpret data by considering all possible factors. There are other mechanisms not explicitly discussed in this paper, such as spatial pattern of cell differentiation, chromatin remodeling and DNA methylation [62–64].

The time-dependent parameter governed by a timescale factor corresponds to the delay activities in gene regulation. Another way of investigating the effect of time delay in cell differentiation (e.g., delay-induced oscillations) is by employing a delay differential equation model (for example, see [65]). Differential equation models with time delay are generally considered as infinite-dimensional systems.

In summary, our simulations predict the following outcomes: First is the sequential branching of lineages in cell-fate determination that portrays differentiation from pluripotent state to transient states (lineage progenitors) towards specialized cell types. Second is the cellular reprogramming driven by oscillations generated by a repressilator-type network, which is a possible technique for dedifferentiation or transdifferentiation. A two-variable switch-like model usually cannot illustrate the branching phenomena and sustained oscillations, but a model with multiple GRFs and asymmetric reciprocal interaction between GRFs can. Oscillatory behavior cannot be taken for granted because this could explain peculiar dynamics related to the epigenetic machinery of organisms, such as dedifferentiation as part of regenerative process. Oscillations are also involved in pattern formation, circadian rhythms, and the progression of diseases [46,49,53,66–70]. Network motifs, such as the repressilators, that are part of a larger GRN induce functional fluctuations necessary for tissue development and cellular engineering. Investigating the dynamics of these network motifs can be helpful in drug discovery [71].

## Methods

In our simulations, we use the following differential equation model (see Box 1):

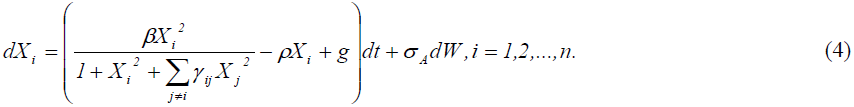

We assume the following parameter values: *β=1, ρ=0.05* and 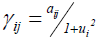. The GRFs have the same value of *β* and *ρ* to highlight the effect of time-varying *γ*_*ij*_. To test the robustness and sensitivity against moderate random noise, we introduce the term *σAdW* representing Gaussian white noise with amplitude *σA.* Let *σA=0* and *σA=0.5* for deterministic and stochastic simulations, respectively. The noise term approximates multiple heterogeneous sources of additive random fluctuations.

### Time evolution of the interaction coefficient

The value of 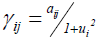 is updated using the following gradient function:

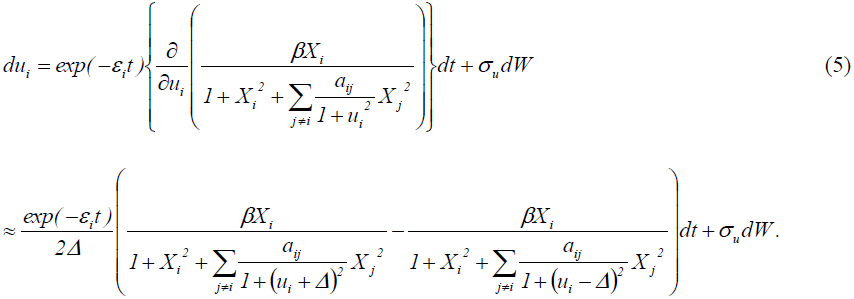

Equation (5) is used for finding relative optimum and is similar to the trait dynamics frequently used in evolutionary biology [72,73]. The trait dynamics in fitness landscape (a concept in evolutionary biology) is similar to the cell-fate determination dynamics in Waddington’s epigenetics landscape [58].

Increasing the value of *u*_*i*_ decreases the value of repression coefficient *γ*_*ij*_ (*j≠i*). Hence, the variable *u*_*i*_ can be defined as the time-varying attribute for maximizing the strength of the GRF *X*_*i*_. However, note that increasing the value of *u*_*i*_ does not always result in an increased equilibrium value of *X*_*i*_, especially when the initial condition and other parameter values do not allow significant changes in the epigenetic landscape in favor of *X*_*i*_.

The timescale factor is represented by *exp*(*-ε*_*i*_*t*) with decline rate *ε*_*i*_, as described in various studies [5,12]. As time progresses (e.g., as cell matures), the timescale factor declines and the value of *u*_*i*_ leads to equilibrium. In addition, the dynamic parameter *u*_*i*_ is initialized with value *u*_*i*_*(0)=0.001* for all *i.* Let *σ*_*u*_=0 and *σ*_*u*_*=0.01* for deterministic and stochastic simulations, respectively. For simplicity, we approximate the partial derivative in Equation (5) using central difference formula with *Δ=0.001.*

### Quasi-potential, cell type probability and network entropy

We suppose that the quasi-potential (*Φ*) of the epigenetic landscape is computed as

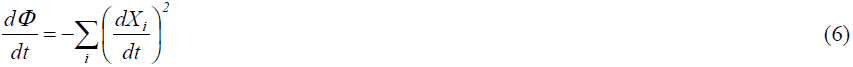

where the 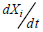 is deterministic [6,17]. This quasi-potential is assumed to characterize the elevation of the landscape [6]. The basal constitutive growth rate *g*, which can also represent effect of external stimulus, is excluded in our computation of the quasi-potential.

The probability of differentiating to cell type *i* or the expected proportion of cells committed towards cell type *i* is 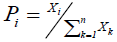. To visualize the canalization in Waddington’s illustration, we use three coordinate axes: time, quasi-potential, and cell type probability. We also compute for the network entropy [19, 74] defined by

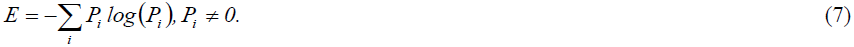

This network entropy measures the potency of population of cells. Population of multipotent and pluripotent cells have high network entropy because they can proliferate to heterogeneous cell types.

### Numerical method

The ordinary and stochastic differential equations are solved using Runge-Kutta 4 and Euler-Maruyama with 0.01 as step size, respectively. For supplementary mathematical discussions about the differential equation model (Box Eq. 1), refer to related literatures [10,18,35].

### Notes on abstraction

One of the advantages of the model in Box 1 is that it is straightforward, thus any peculiar dynamics can be clearly interpreted. However, the model is an abstraction of the cell differentiation process. Hence, we focus on the qualitative dynamics rather than on exact quantitative values. Likewise, the qualitative dynamics arising from the model are possible to arise in a more complex system that contains the minimal GRN (Fig. 1b) as a subnetwork.

## Acknowledgment

We would like to thank B.D. Aguda and C.O. Talaue for discussion about models of cellular regulation, and the organizers and participants of the 2014 Hands-on Research in Complex Systems School at ICTP for feedback and suggestions. We would also like to thank E.R. Mendoza, R.C.H. del Rosario and our colleagues from the mathematical and biological sciences whom we have requested to review our manuscript. JRF is supported by Japanese government (Monbukagakusho: MEXT) fellowship and the University of the Philippines. ALB is supported by Philippine Commission on Higher Education (CHED) fellowship and the University of the Philippines.

## Author contributions

JFR and ALB conceived the study. JFR built the model and ran the simulations. JFR and ALB wrote the manuscript.

## Supplementary Materials

Supplementary Figs. 1 to 21

